# Emergence of Directional Actomyosin Flows from Active Matter Vibrations

**DOI:** 10.1101/394700

**Authors:** Sven K. Vogel, Christian Wölfer, Diego A. Ramirez-Diaz, Robert J. Flassig, Kai Sundmacher, Petra Schwille

## Abstract

Cortical actomyosin flows play pivotal roles in cell motility, cell division and animal morphogenesis. According to many model systems, myosin motor induced local contractions are key for generating cortical flows. However, the original mechanism how large-scale directed flows emerge from local motor activity in an apparently isotropic cortex is unknown. We reconstituted and confined minimal actomyosin cortices to the interfaces of emulsion droplets. The presence of ATP leads to myosin-induced cortical contractions that self-organize into directed flow-like actomyosin motions. By combining our experiments with theory, we found that the large-scale directional motion of actomyosin clusters emerges from individual asymmetric cluster vibrations, caused by intrinsic non-isotropic ATP consumption, in conjunction with spherical confinement. By tracking individual actomyosin clusters, we identified fingerprints of vibrational states as the basis of directed motions. These vibrations may represent a generic key driver of directed actomyosin flows under spatial confinement *in vitro* and in living systems.

---

In animal cells, cortical actomyosin motions including actomyosin flows have been proposed to drive cell locomotion, cytokinesis, left-right symmetry breaking during embryonic development of multicellular organisms, and cellular or tissue chirality (*1-6*). Despite the omnipresent functions and implications of cortical actomyosin motions and flows, the exact molecular origins and fundamental determinants of these phenomena are far from being understood. In many eukaryotic model systems, myosin motors acting on actin filaments are proposed to play a key role in driving actomyosin dynamics in cytokinetic rings and cortical actomyosin flows (*1, 4, 7-9*). Distinct manipulation of myosin and actin independent of other cellular processes is challenging, since they both are functionally highly integrated cellular proteins. Moreover, to elucidate whether directional flows may result spontaneously from active processes without being guided by the structural anisotropy of the cellular architecture, it is mandatory to explore these phenomena in well-controlled reconstituted systems (*10, 11*). Recent studies used the approach of encapsulating frog egg cell extracts in droplets where manipulation of the protein players may be easier to achieve compared to living systems (*12, 13*). Nevertheless, due to the compositional complexity of cell extracts comparable to living model systems, it may be difficult to pinpoint the minimal set of necessary proteins to produce a certain phenomenon. High density regimes of nematic cytoskeletal *in vitro* systems showed large-scale pattern formation and dynamics in a collective manner based on motor driven sliding mechanisms between long cytoskeletal filaments (*14, 15*). In contrast, actomyosin cortices and rings in living systems consist of non-polar and disordered actin filament networks and are coupled to the cell membrane (*16, 17*). By a combination of bottom-up *in vitro* experiments and theory, we identified a generic mechanism of how large-scale directional flow-like actomyosin motions are generated by myosin motors in a disordered, membrane coupled and isotropic cytoskeletal system in a confined spherical environment. To this end, we made use of recently developed minimal actin cortices (MACs) (*18, 19*) and confined them in water-in-oil droplets using microfluidic emulsification on PDMS chips. The chip design is shown in Fig. 1A. (Note that we also performed emulsification simply by manually mixing the components to rule out any suspicion of flows caused by the pneumatic microfluidic setup.) DOPC and biotinylated lipids were dissolved in mineral oil, and for droplet formation we used a pneumatic microfluidic system where the flow rate of each channel is individually controlled (Movie S1). The aqueous solution contained dissolved actin monomers, neutravidin and myosin in a salt buffer system. To test the proper formation of a lipid monolayer that includes biotinylated lipids (DSPE-PEG-2000-Biotin) we started with the encapsulation of fluorescently labeled neutravidin (Oregon Green 488 Neutravidin) (Fig. 1B-D). By analyzing the fluorescence intensity through the droplets, we showed that neutravidin only binds to the lipid monolayer when the lipid oil mixture also contains biotinylated lipids (Fig. 1 C and D) indicating that neutravidin does not bind non-specifically to the water oil interface. Co-encapsulation of myosin motors with the actin monomer and anchor system in the presence of ATP resulted in the formation of a MAC. Inside the droplets, actin monomers start to polymerize and bind to the lipid monolayer that contains biotinylated lipids (Figs. 1B and E, Movie S2). Concurrently, ATP-induced contraction of the myosin motors resulted in the formation of actomyosin clusters (Fig. 2A-C, Movie S3). Strikingly, within minutes a large-cale directional movement emerges from these spherically confined actomyosin clusters which we will refer to as cortical actomyosin motion (CAM) (Figs. 1E and 2B; Movies S3 and S5). The CAMs can last more than one hour before they eventually cease, reaching a so-called frozen state with only minimal movements (Movies S3 and S5). Interestingly, these flow-like motions somewhat resemble cortical actomyosin flow-like motions *in vivo*, e.g. in the *C. elegans* embryo. We characterized the velocity of the actomyosin cluster movements by using particle image velocimetry (*20*) (Fig. 2C; Movie S4). In order to test parameters that change the velocity of CAMs, we co-encapsulated a crowding agent (methylcellulose) at various concentrations, and found a respective increase of the velocities in the presence of methylcellulose (Fig. 2D, compare Movies S3 and S5). By increasing the effective concentration of actin and myosin at the lipid monolayer membrane interface, the overall cluster velocity was increased to more than five times.

**Fig. 1.**
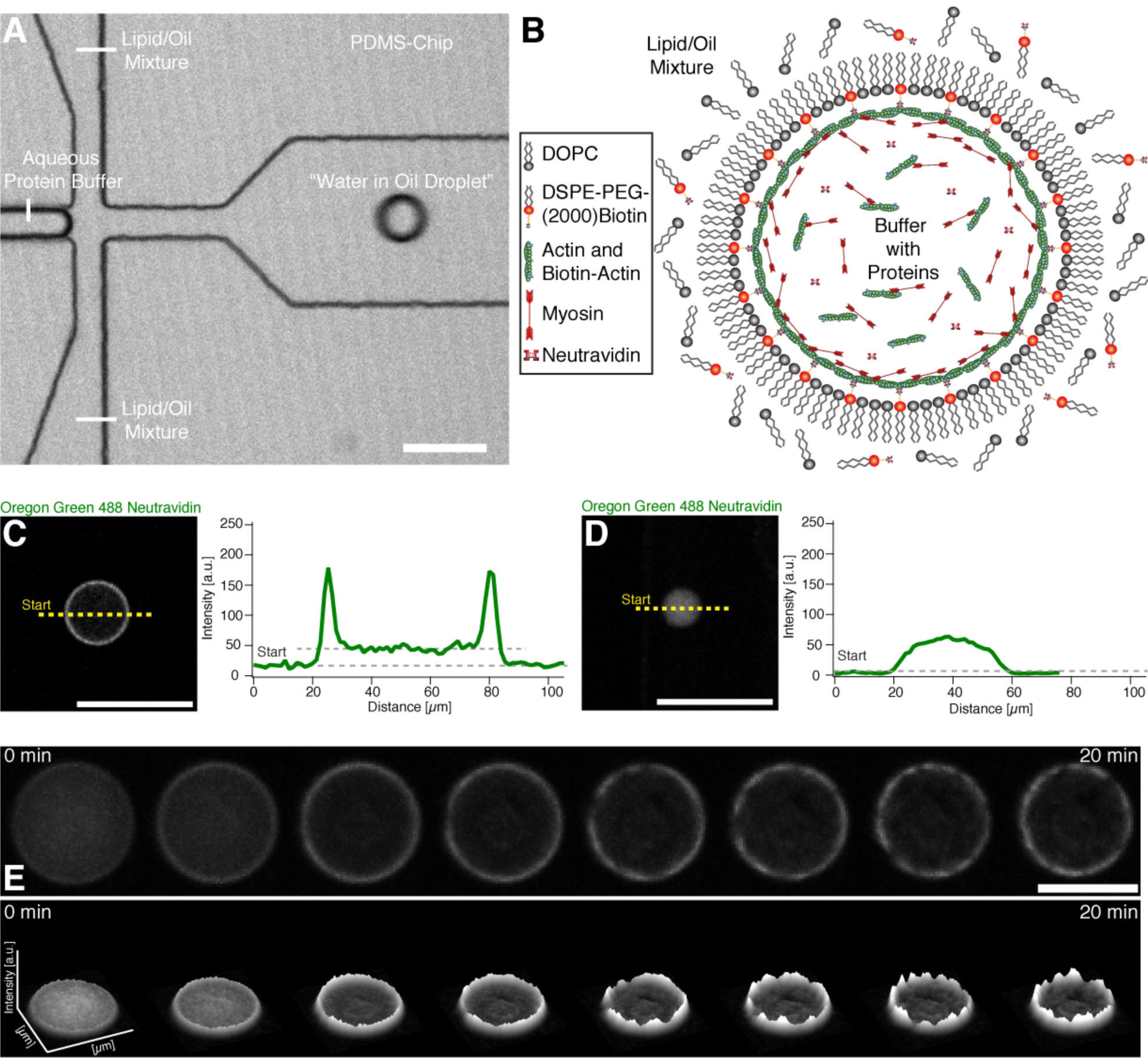
Encapsulation and actin cortex formation inside water in oil droplets. (A) Confocal image of a PDMS chip where the encapsulation of the buffer-protein system (see (B)) and formation of the water in oil droplet is shown (Movie S1). (B) Illustration depicting the formation of an actomyosin cortex. (C) Confocal image of the equatorial plane of a droplet with a lipid monolayer containing biotinylated lipids showing that encapsulated Oregon green labeled neutravidin binds to the lipid monolayer. Line profile of the fluorescence signal of the Oregon green labeled neutravidin shows two peaks which indicate binding of the neutravidin to the lipid monolayer interface of the droplet (right). (D) In contrast Oregon green labeled neutravidin does not bind to the lipid monolayer and is distributed throughout the lumen of the droplet in the absence of biotinylated lipids. The fluorescence line profile shows no peaks in the absence of biotinylated lipids indicating the absence of unspecific binding to the lipid monolayer (right). (E) Confocal time-lapse images of encapsulated Alexa-488 labeled actin and myosin motors in the presence of ATP at the droplet equator plane. The formation of an actin cortex and actomyosin clusters is shown (upper row) (Movie S2). The respective fluorescent intensity profile indicates the formation of actomyosin clusters and shows their dynamics (lower row). Scale bars, 100 µm.

**Fig. 2.**
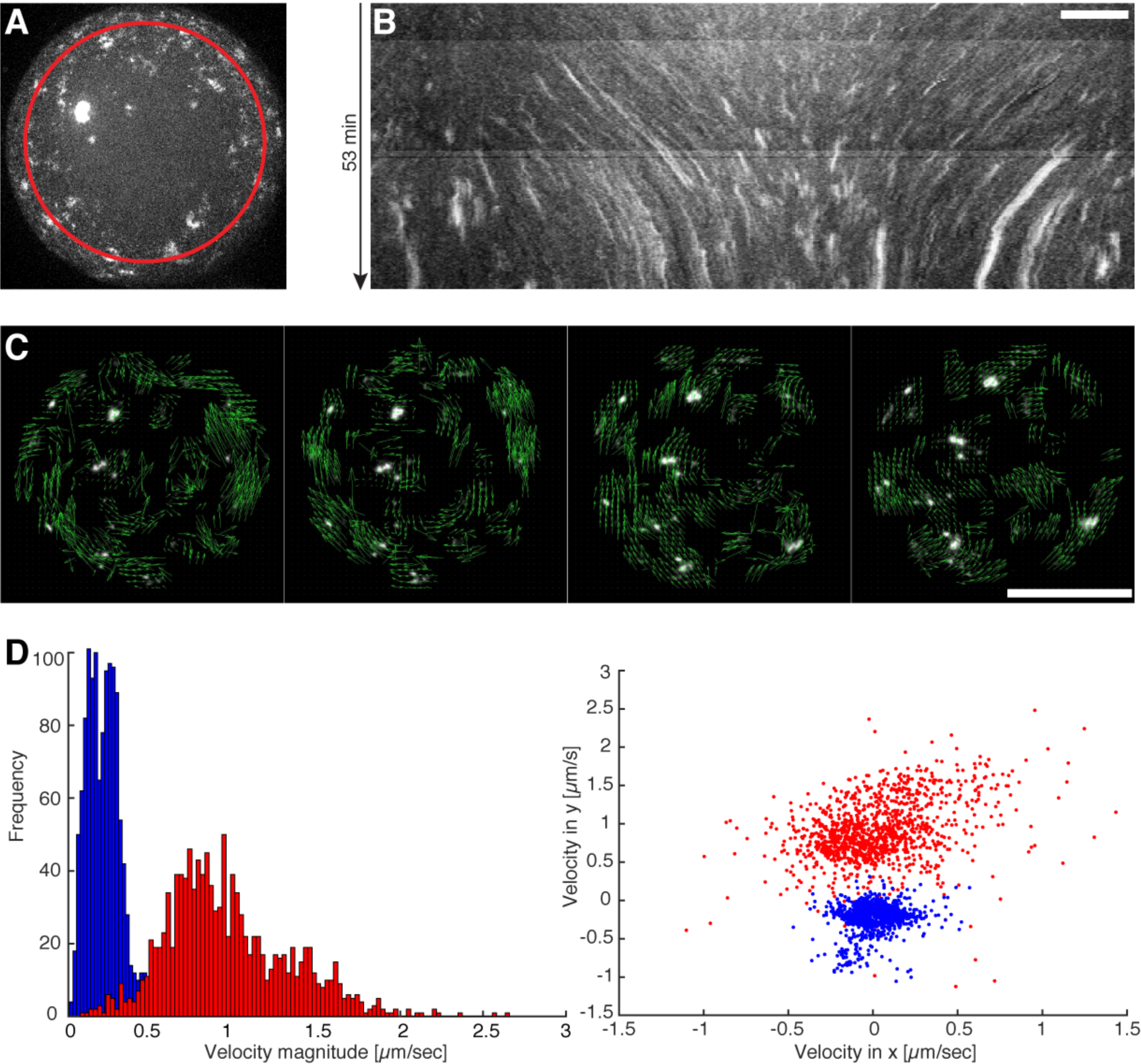
Directed movement of actomyosin clusters upon ATP dependent actomyosin contractions. (A) Maximum intensity projection from a half droplet confocal z-stack where Alexa-488 labeled actin clusters are visible (Movie S3). The red circle indicates the path of the generated kymograph. (B) A kymograph of the maximum intensity projected half sphere is shown where directed movements of individual clusters represented by distinct lines are visible. Scale bar, 10 µm. (C) Confocal time-lapse image sequence used for particle image velocimetry (PIV) by (*20*). Vectors (green arrows) indicate the flow direction of the directed movement of the actomyosin clusters (Movie S4). Scale bar, 50 µm. (D) Two examples of velocity profiles measured by PIV of droplets with (red) and without a crowding agent (blue) are shown (Movies S3 and S5).

We now aimed to elucidate the mechanism behind myosin driven CAMs, and complemented our experiments with a theoretical model based on first biophysical principles, i.e. without assuming an explicit mechanism *a priori*. We modeled the MAC droplet as a continuous, isotropic fluid system. The interaction between actin and myosin was modeled by a simplified myosin cross-bridge cycle (21-23) where the force generating conformational change (r_1_) of the myosin head (M) occurs immediately after the binding of filamentous actin (A) (Fig. 3A). The actomyosin complex dissociates after binding ATP whose hydrolysis reloads the myosin head into the active state (r_2_). Usually considered intermediate species have only a very short lifetime (*24*) and are therefore neglected. As the MACs were formed by polymerization, in the absence of actin-regulating proteins (*25-27*), a rudimentary polymerization cycle (*24*) was added to the model. F-actin depolymerizes (r_3_) into monomeric G-Actin with bound ADP (G_D_), followed by a spontaneous nucleotide exchange (r_4_). The resulting ATP bound G-Actin (G_T_) binds stronger to F-actin (r_5_) than GD-actin (r_6_), and therefore GT-actin polymerization is dominant (28). F-actin, cross-linked by myosin II molecules, forms a mesh-like cortex causing an active viscoelastic material behavior with a rheological property combination of the Maxwell and Kelvin-Voigt models (*29*) considered in the momentum equation (*30*) (Supplement Eq. S14). To model force generation according to the myosin cross-bridge sub-model, the active stress was developed from a stress term of our previous study (*31*) considering the observed medium ATP-dependency of myosin pulls in MACs (*19*). The distributed reaction network is described by a system of partial differential equations (PDEs) assuming diffusion (G_D_, G_T_, ATP) or combined convection and diffusion flux (A, A_M_, M) (Supplement Eq. S1-S6). We reduced our analysis to a one-dimensional ring topology modeling a section of the spherical droplet, to improve and simplify the interpretation of the simulation results. To adopt experimental conditions, the PDE system was simulated with an initially slightly inhomogeneous myosin distribution and revealed merging of small clusters to a gradually bigger main cluster similar to experimental observations (Fig. 3B and C). The final non-symmetric cluster starts to move directionally in the one-dimensional ring system like a propagating wave (Fig. 3B and C). Hereinafter, two characteristics are described, which are presumably essential for this movement. First, a non-symmetric contraction profile is preserved by an unbalanced consumption of ATP at the leading and trailing edge of the propagating cluster. Because of the locomotion and the ongoing network depolymerization, the cluster traces non-convective GD-actin like a comet tail (fig. S1). In the GD-actin tail, ATP is additionally consumed for the nucleotide exchange of monomeric actin (r_4_). Thus, less ATP is available for the force-generating myosin cross-bridge cycle at the trailing edge of the wave and hence less contractile stress is generated compared to the leading edge (Fig. 3E). Second, examining the fine structure of the simulated moving cluster reveals a vibration. It is driven by periodical depletion of the local ATP concentration, leading to oscillating active and passive stresses inside the cluster (fig. S3A). The cluster vibration passes through the following repetitive phases (Fig. 3F):

I. *Contraction Phase*: In the beginning of the contraction phase, a high ATP concentration inside the cluster results in an increase of the contractile stress accompanied by an increase of elastic stress. With progressing cluster compaction, the actomyosin species gets locally accumulated, causing stronger contractile stress and increased local ATP consumption.
II. *Expansion Phase*: Increasing local ATP depletion and network strain leads to a local dominance of elastic stress resulting in an expansion of the cluster. Because of the reduced network densities inside of the cluster and therefore reduced energy consumption, ATP flows in diffusively and increases the local ATP level, followed by a de-novo contraction phase.

**Fig. 3.**
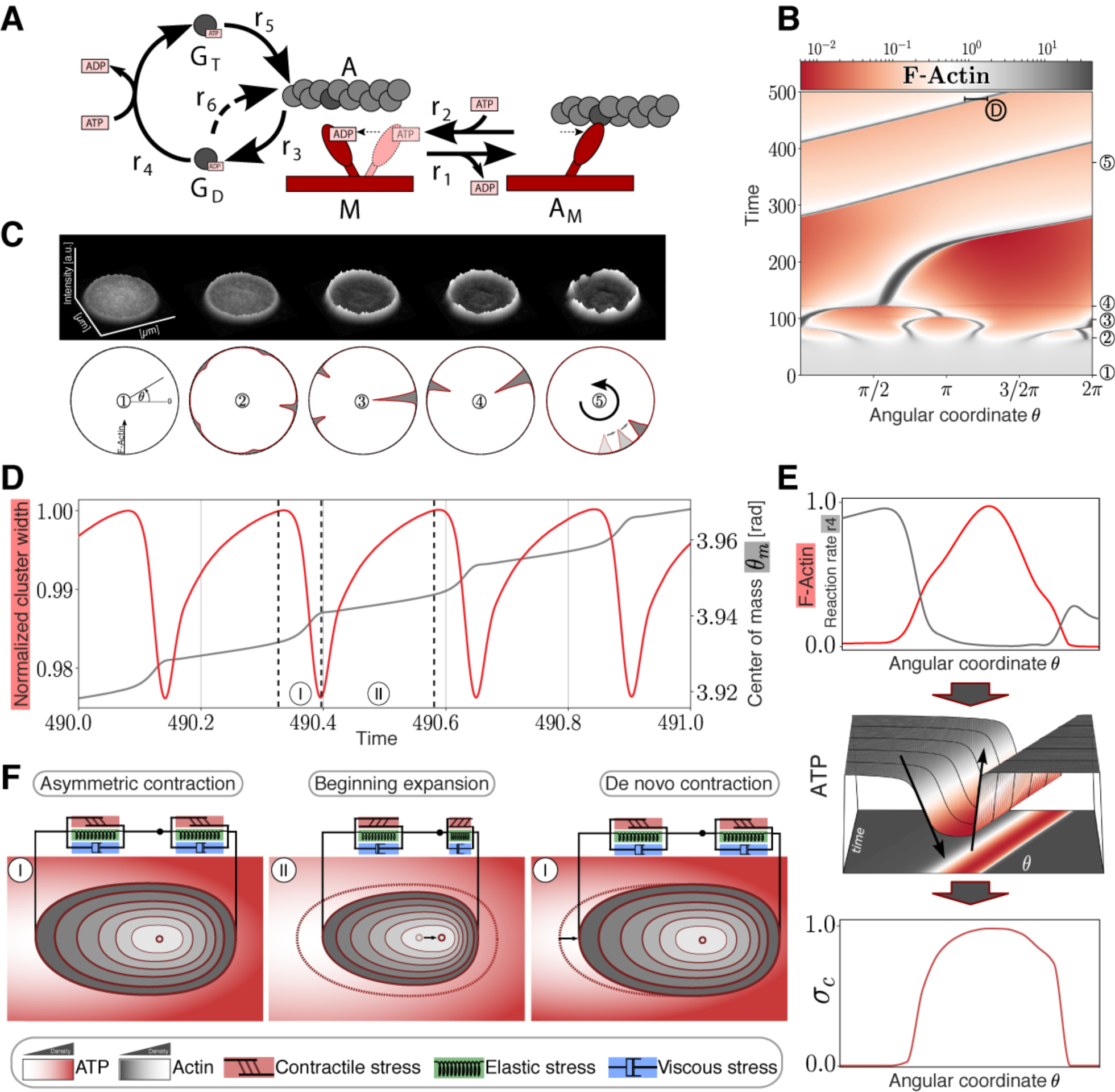
Modeling and simulation of actomyosin cluster motions reveal a propagation mechanism through active matter vibrations. (A) Kinetic reaction network with F-actin polymerization cycle and simplified myosin cross-bridge model. (B) Temporal development of a one-dimensional spatial F-actin distribution with color-coded local concentration. (C) Circular representation of distributed F-actin concentration according to the model topology for selected time points (marked in 3B with circles). A fluorescent intensity profile of the formation of actomyosin clusters and their dynamics from experimental data (see also Fig. 1E) is shown for comparison (upper row) (D) Vibration of an F-actin cluster. Evolution of normalized cluster width (red) and displacement of center of mass *θ_m_* (gray) of F-actin in radiant. Cluster boundaries are defined as the points where the F-actin concentration exceeds the mean concentration. (E) Non-symmetric contraction of a propagating cluster. Upper graph: Normalized distribution of reaction rate 4 (gray) compared to normalized F-actin cluster location (red). Middle graph: Non-symmetric ATP distribution around the cluster. Lower graph: Normalized asymmetric contractile stress pattern. (F) Sketch of the cortical actin cluster migration mechanism with qualitative color-coded Actin and ATP gradients shows repetitive asymmetric contractions (I) (I) and an expansion phase (II) resulting in a shift of the center of mass indicated by a black arrow (middle panel) and displacement of the cluster indicated by a black arrow (right panel).

In addition, the non-symmetric contraction pattern causes an accelerated displacement of the center of mass during the contraction phase (Fig. 3D and F). Owing to viscoelastic material properties and asymmetric creep, the center of mass is not pushed back to the initial position during the expansion phase, as a result of the non-symmetric contraction pattern. Since the active stress is ATP-driven, the wave propagation ceases owing to the global ATP depletion (fig. S2). For further investigation of the wave propagation mechanism, we assume a constant diffusive ATP supply. Hence, cluster vibrations due to local ATP depletion and asymmetric cluster contractions are the main drivers of single cluster motions. In contrast to previous theoretical studies (*32, 33*), where the required asymmetry of a self-propagating cluster is caused by polar actin bundles, our suggested model is also able to explain cluster migrations of disordered actin filaments that can be expected in an isotropic cortex.

To find experimental evidence for the theoretically predicted vibrations of the actomyosin clusters, we automatically tracked their directional movement (Fig. 4A). In addition, by analyzing the directed movement of the individual actomyosin clusters, we noticed that rotation of clusters around their center of mass, correlated with a change in direction displacement (Fig. 4A and B; Movies S6 and S7). This can be explained with torque produced by an imbalance of forces perpendicular to the translational trajectory. On the other hand, the model predicts vibrations with a certain oscillation period (Fig. 3D). Evidence for such vibrations in our experimental conditions can be found independent of the sampling rate. For example, a sinusoidal signal with a specific frequency can be measured at undersampling conditions (fig. S5). The reconstructed signals still conserve the periodic behavior reflected by the Fourier analysis, yet not reflecting the original frequency (fig. S5). To find evidence for vibrations inside the clusters as predicted by the theory, we analyzed image sequences to measure fluctuations in the distance between the geometrical and the intensity-weighted center of mass of several clusters. The geometrical center of mass was computed by regular segmentation and binarization defining the geometry of the cluster. In contrast, the intensity-weighted center of mass was calculated over the same geometry but having the camera-intensity as the statistical weight of every pixel. Since clusters have various geometries, we defined a characteristic length for the clusters as the square root of the area. This quantity allowed us to measure changes between the geometrical and intensity centers of mass with respect to their characteristic length. As a result, we were able to combine fluctuations of myosin clusters with different sizes and experimental conditions (fig. S6). Fourier analysis of these combined fluctuations strikingly indicates the existence of vibrational states (Fig. 4D, red, blue and yellow curves) which are clearly distinguishable from no-movement conditions (Fig. 4D, grey curve) or acquisition and camera artifacts by measuring the background between moving clusters (data not shown as the curve is similar to the static conditions). We interpret these vibrational states as fingerprints of the theoretically predicted vibrations that provide a mechanism for the directed and rotational large-scale movements of the actomyosin clusters *in silico* and *in vitro*. Under crowding conditions where CAMs show the highest velocities, we found higher amplitudes for the same observed frequencies in comparison to the other tested conditions (Fig. 4D). This finding implies that larger amplitudes correlate with higher velocities, as expected for vibrations-based movements. Since the crowder increases the effective concentration of actin and myosin at the lipid monolayer membrane interface, we conclude that higher actomyosin concentrations may lead to higher cluster velocities and therefore to larger amplitudes of the vibrations.

**Fig. 4.**
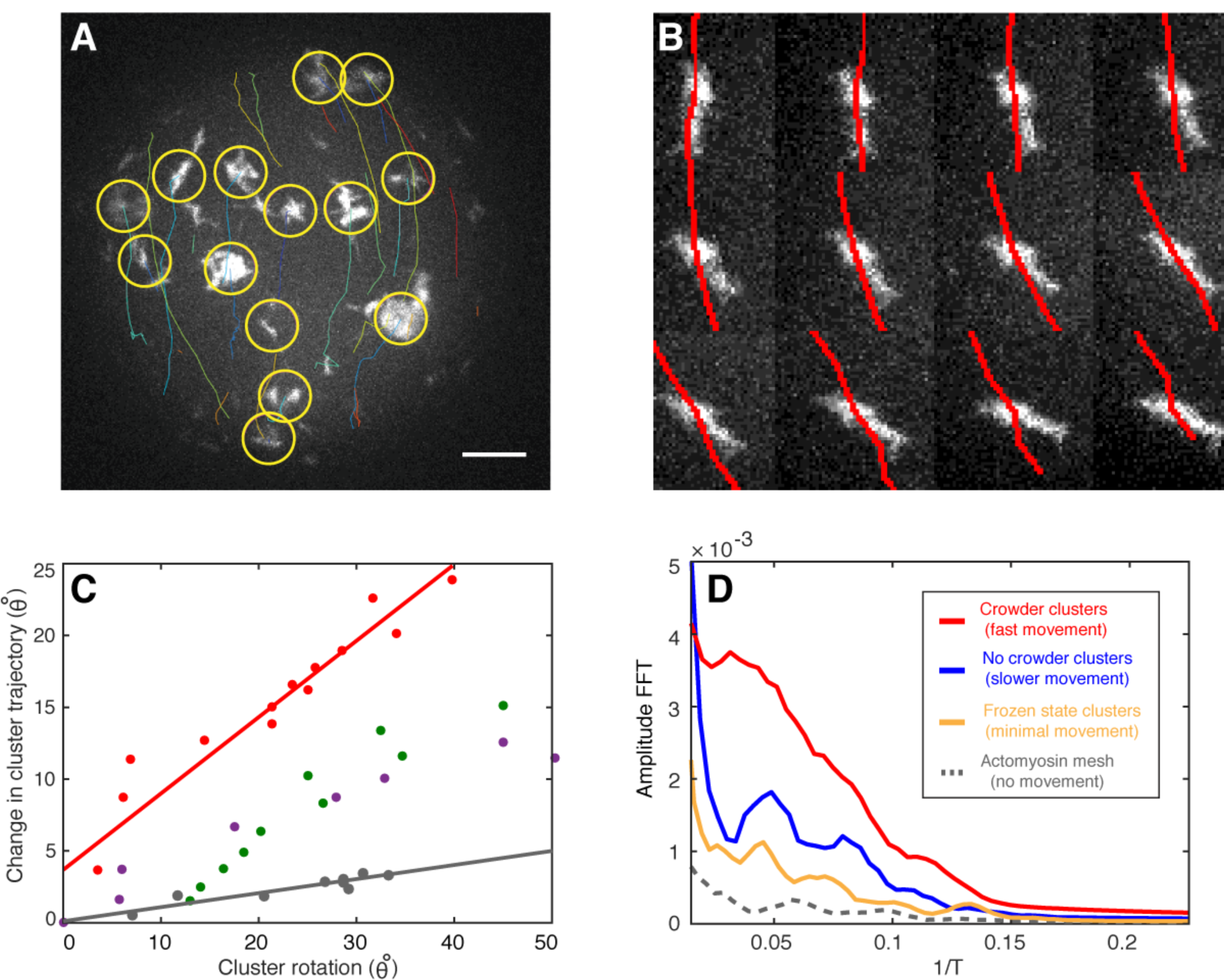
Fingerprints of actomyosin cluster vibrations during their directed movements. (A) A maximum intensity projection of confocal images of a half sphere during cortical actomyosin cluster movements is shown. Tracked actin clusters are marked by circles (yellow) and their trajectories shown (various colors, Movie S6). (B) The rotation of clusters along their center of mass agrees with changes in the trajectory (red lines) direction (left, Movie S7). (C) Rotation of four independent clusters (various colors) show a correlation between cluster rotation and different rates of steering (different slopes, red and grey line). The red data points correspond to the montage shown in B. (D) The fast Fourier transformation (FFT) indicates the presence of vibrational states for moving actomyosin clusters (red, blue and yellow) different from static systems and acquisition artifacts (grey). The higher amplitude for the crowder condition with the highest cluster velocities of all measured systems implies that larger amplitudes correlate with higher velocities. Scale bar, 10 µm.

Based on our suggested mechanism for the directional motion of an individual cluster, we consider the possibility of an alignment of motion of a group of clusters in one direction, resulting from potential cluster interactions via the cortical actin network, thereby explaining the observed flow-like group behavior of the actomyosin clusters. Alternatively, a macroscopically homogeneous actin carpet is initially formed as shown in our experiments (see Fig. 1E, Movies S2 and S3). Since the resulting clusters are initially connected via the actin mesh, the random asymmetry of the largest cluster would determine the direction of movement of the entire cluster group, in the absence of coordinating processes. Each cluster of the group is thus impressed with a direction of movement or an asymmetric gradient profile originating from the respective monomer tail. In conclusion, our theory gives evidence that the direct translational and rotational movement of actomyosin clusters originates from an imbalance of oscillatory contractile stresses within the individual actomyosin clusters. Fourier analysis of our experimental data indicates the existence of vibrational states that drive the directional movements of individual actomyosin clusters and the formation of flow-like CAMs in the spherical confinement of the active matter droplets. Hence, for future work it will be of great interest to investigate whether these vibrational states can be also identified in cellular model systems that perform myosin driven actomyosin motions, e.g. in cytokinetic actomyosin rings or in cortical actomyosin flows.

## Acknowledgements

We are grateful for the financial support by the MaxSynBio consortium jointly funded by the Federal Ministry of Education and Research of Germany and the Max Planck Society, the Daimler und Benz foundation (Project Grant PSBioc 8216), the Gottfried Wilhelm Leibniz-Program of the DFG (SCHW 716/8-1) and the support of the Graduate School of Quantitative Biosciences Munich.

